# siRNA Features - Reproducible Structure-Based Chemical Features for Off-Target Prediction

**DOI:** 10.1101/2025.03.24.645065

**Authors:** Michael Richter, Alem Admasu

## Abstract

Chemical modifications are the standard for small interfering RNAs (siRNAs) in therapeutic applications, but predicting their off-target effects remains a significant challenge. Current approaches often rely on sequence-based encodings, which fail to fully capture structural and protein–RNA interaction details critical for off-target prediction. In this study, we developed a framework to generate reproducible structure-based chemical features, incorporating both molecular fingerprints and computationally derived siRNA–AGO2 complex structures. Using an RNA-Seq off-target study, we generated over 30,000 siRNA–gene data points and systematically compared nine distinct types of feature representation strategies. Among the datasets, the highest predictive performance was achieved by Dataset 3, which used extended connectivity fingerprints (ECFPs) to encode siRNA and mRNA features. An energy-minimized dataset (7R), representing siRNA–AGO2 structural alignments, was the second-best performer, underscoring the value of incorporating reproducible structural information into feature engineering. Our findings demonstrate that combining detailed structural representations with sequence-based features enables the generation of robust, reproducible chemical features for machine learning models, offering a promising path forward for off-target prediction and siRNA therapeutic design.

## Introduction

Despite the growing interest in leveraging machine learning and computational techniques for chemical data and siRNA therapeutics, a comprehensive exploration of feature engineering in this domain remains significantly underdeveloped. A thorough review of recent literature (2023–present) using advanced search strategies in Web of Science (WOS) yielded only a single relevant review article. This scarcity of resources highlights the novelty and unexplored nature of applying feature engineering and machine learning to chemically modified siRNA design and off-target prediction. The current gap underscores the need for new studies to establish foundational frameworks and methodologies in this critical intersection of cheminformatics, RNA therapeutics, and machine learning. Small interfering RNAs (siRNAs) are short, double-stranded RNA molecules that use the RNA interference (RNAi) pathway to silence gene expression post-transcription. By guiding the RNA-induced silencing complex (RISC) to complementary mRNAs, siRNAs induce targeted degradation, effectively preventing production of specific proteins. Owing to their precision and broad applicability, siRNAs hold promise for treating a variety of conditions, including cancer, genetic disorders, and viral infections.

A major challenge in siRNA therapeutics is the unintended off-target effects, where siRNAs bind to partially complementary mRNA sequences or interact with unintended proteins. These off-target interactions can lead to undesirable gene silencing, toxicity, and immune activation, undermining therapeutic efficacy and safety. The seed region of the siRNA guide strand (positions 2–8) is particularly prone to off-target binding, as it drives initial mRNA recognition. To address these issues, chemical modifications have emerged as powerful tools to enhance siRNA specificity and reduce off-target interactions, and they are now considered the standard in modern siRNA therapeutic design.

The discovery of effective chemical modifications for siRNA faces several challenges. These include balancing stability, efficacy, and safety, particularly in minimizing off-target effects. Chemical modifications must preserve RNAi activity while reducing off-target interactions. However, understanding how specific modifications impact the complex interplay of siRNA–protein and siRNA–mRNA interactions remains limited. Advances in computational modeling and high-throughput experimental platforms provide new opportunities to explore novel chemical structures and predict their biological outcomes. Nevertheless, because these chemical modifications often involve noncanonical or “unnatural” structural elements, conventional sequence-based off-target prediction methods cannot adequately capture them, thereby reinforcing the need for new approaches. Integrating these approaches is critical for optimizing siRNA therapeutics to address unmet clinical needs.^1^

As this is an emerging field, a review of recent preprints was conducted to capture the latest innovations in siRNA off-target prediction, feature engineering, and structure-based modeling before formal peer-reviewed publication. Several significant works have appeared within the last year, covering a range of topics, including machine learning approaches, molecular modeling, bioinformatics tools, and structural biology insights.

Oshunyinka introduced a machine-learning framework using sequence-derived features to predict siRNA potency.^2^ His model, tested against SARS-CoV-2, underscoring how careful encoding of siRNA sequences can capture structure– potency relationships. Similarly, Bai et al. presented OligoFormer, a transformer-based deep learning model that jointly predicts siRNA knockdown efficacy and off-target potential.^3^ The model incorporates thermodynamic parameters and pretrained RNA embeddings, and Bai et al. further validated it using microarray data and integrated TargetScan and PITA to refine off-target assessments. Long et al. proposed a GNN-based approach (published as siRNADiscovery) that represents siRNA–target pairs as a bipartite graph, leveraging message-passing to capture long-range dependencies in sequence context. While that study focused on efficacy, the authors noted its potential extension to modeling off-target interactions.^4^

Recent advances in feature engineering underscore the importance of chemical and sequence-based featurization. In a pair of complementary approaches, Liu et al. introduced Cm-siRPred—a machine learning model that integrates ECFP-based molecular fingerprints, k-mer encoding, and target site accessibility metrics for chemically modified siRNAs—and AttSiOff, a self-attention-based model coupling efficacy prediction with an off-target filtration module that ranks 3′UTR sequences by seed matches.^5^ Yadav et al. developed a stochastic model of siRNA endosomal escape, showing how incomplete cytosolic release can limit effective siRNA concentration and potentially increase off-target effects at high doses.^6^ Cazares et al. released SeedMatchR, an R package that annotates RNA-seq data from knockdown experiments to flag differentially expressed genes harboring 6–8mer seed matches, providing an open-source workflow for off-target analysis.^7^

Beyond machine learning, several structural biology preprints have yielded important insights into siRNA–AGO2 interactions. Sarkar et al. reported a 3.16 Å cryo-EM structure of human Argonaute2 bound to a guide siRNA and target mRNA, revealing a distortion at position 6 that repositions Lysine-709 in the catalytic site—a previously unobserved mechanism.^8^ Wallmann and Van de Pette conducted a comparative evolutionary analysis of Argonaute, identifying conserved atomic interactions that stabilize the siRNA guide-strand and suggesting that future models incorporate protein-contact features.^9^ Finally, Enoki et al. combined explainable AI with wet-lab validation to discover a novel RNA-binding protein that modulates siRNA selective loading into RISC complexes, highlighting AI-guided bio-discovery’s potential for advancing RNAi research.^10^

Collectively, these preprints reinforce the pivotal role of structural insights, feature engineering, and data-driven models in refining siRNA design, emphasizing the growing reliance on sophisticated machine learning approaches to tackle off-target complexities.

Machine learning (ML) has become indispensable for siRNA efficacy and off-target prediction, as it can uncover complex patterns beyond rule-based design. However, the quality of input features is critical to model success.^11^ Traditional sequence encodings (e.g., one-hot vectors or positional indices) and simple chemical descriptors (e.g., SMILES strings) often fall short, as they primarily capture linear information and overlook spatial or interaction context crucial for RNA–protein complexes.^11,12^ Recent studies have explored advanced feature engineering to address this gap. For instance, graph neural networks (GNNs) show promise in small-molecule tasks but face scalability challenges with large biomolecular assemblies like siRNA–Argonaute complexes.^13^ Current GNN frameworks (e.g., DGL) are not optimized for the size and structural complexity of protein–RNA interactions, limiting their effectiveness for siRNA off-target modeling. This has motivated the development of alternative feature representations that can capture three-dimensional and chemical nuances of siRNA molecules.

Researchers are increasingly combining sequence-based features with structure-based and physicochemical features. Platforms like iFeatureOmega integrate diverse encodings – from k-mer frequencies and position-specific scoring matrices to biochemical property profiles – yielding richer representations of RNA sequences.^11^ Such integrative features can improve model performance but also risk overfitting, especially with limited training data. Therefore, a balance between feature richness and generalizability is essential.

To represent chemical modifications in siRNAs, cheminformatics tools offer powerful solutions. Extended-connectivity fingerprints (ECFPs), for example, encode molecular substructures into fixed-length vectors.^14^ Unlike NMR-derived fingerprints that require physical compounds,^15^ ECFPs can be computed in silico to capture local atomic neighborhoods of modified nucleotides. In our work, we employ ECFPs – to our knowledge, a novel application in the siRNA context – which effectively encode both standard and chemically modified nucleotides. Early results indicated that compressed ECFP features achieved performance comparable to explicit SMILES embeddings.^16^ This demonstrates ECFP’s potential as a robust, information-rich descriptor for siRNA chemical diversity.

Innovative featurization approaches continue to emerge. One method converts 3D biomolecular structures into 2D images for convolutional neural networks (CNNs).^17^ By projecting protein 3D coordinates (or RNA–protein complexes) into image channels, CNNs can learn spatial features relevant to binding interactions – essentially treating structure prediction as an image recognition problem. Another strategy focuses on thermodynamic stability features: calculating melting temperatures for different regions of the siRNA duplex. Kobayashi et al. showed that the stabilities of the seed region (positions 2–8) versus the 3′ non-seed region (e.g., positions 9–14) have opposite correlations with off-target silencing.^18^ Incorporating such features, including interaction terms combining seed and non-seed stability, provides a biologically interpretable way to enhance off-target prediction.

All these efforts underscore a common theme: spatial and interaction-specific features are key to improving ML models for siRNA. Conventional encodings that ignore 3D context – a limitation noted in analogous domains like chromatin interaction prediction^12^ – can miss critical determinants of binding affinity and specificity. By integrating structural data (e.g., distances in an siRNA– Argonaute complex) and interaction-aware features (e.g., seed pairing stability), we aim to bridge this gap and boost model generalization.

Argonaute proteins are at the core of the RNAi pathway, binding siRNAs and guiding them to complementary mRNAs for cleavage. The guide strand’s 5′ end docks into a hydrophilic pocket formed at the interface of the MID and PIWI domains, anchoring the RNA and positioning it for target recognition. This structural organization ensures precise positioning of the siRNA guide for accurate gene silencing. However, only guide nucleotides g2–g4 are fully exposed and available for initiating interactions with target RNAs, while the remaining guide nucleotides are initially shielded. This pre-organized seed region facilitates rapid and efficient target recognition by lowering the thermodynamic cost of hybridization. Furthermore, the central cleft of Argonaute accommodates the siRNA in an extended conformation, with almost all interactions with the protein mediated through the RNA sugar-phosphate backbone.^19^ While this is true, every atom of the siRNA – from the nucleobases to the backbone – contributes to its overall structural stability, dynamic behavior, and ability to mediate precise interactions. This underscores the importance of incorporating atomic-level detail when modeling siRNA– Argonaute interactions to fully understand their functional impact.

Molecular modeling, especially energy minimization, offers a realistic view of siRNA–Argonaute interactions by detailing spatial arrangements and energetic considerations. Translating these characteristics into ML features can enhance predictive power, potentially improving both efficacy and safety in siRNA therapies. This approach parallels advancements in CRISPR–Cas9 research, where integrating sequence-based features (e.g., nucleotide motifs and positional importance) with non-sequence data (e.g., structural properties, chromatin accessibility, and target site epigenetics) has significantly improved gRNA activity predictions.^20^ By adopting a similarly holistic strategy, combining spatial, energetic, and sequence-based features, siRNA predictive models can achieve greater biological relevance and precision. We hypothesize that the structural features within AGO2, including guide strand docking, seed region exposure, and interactions mediated by the sugar-phosphate backbone, all contribute to the specificity and efficiency of RNAi. Capturing these features at an atomic level and integrating them into machine learning models will enable more accurate predictions of siRNA efficacy and off-target effects, ultimately improving therapeutic design.

We introduce a feature generation strategy that incorporates Argonaute interactions via structural modeling. This proof-of-concept explores whether modeling-derived features can match or exceed conventional approaches, thereby uniting structural biology and ML for more biologically relevant siRNA predictions. Building on current approaches, we developed a novel approach to enhance siRNA feature representation by incorporating 3D spatial information through molecular modeling and simulation. This methodology aimed to address the limitations of conventional features by integrating chemically modified siRNA structures and their dynamic behaviors into predictive models.

To explore off-target interactions, we conducted an RNA-Seq-based study that captured a comprehensive array of off-target effects. From this dataset, the top 20% of genes with the highest mean expression counts were selected for further analysis. Guide siRNA sequences were aligned to each gene to identify intended targets. Using these targets, OpenAI’s Chai Discovery platform^21^ was employed to generate 10,000 structural predictions, forming the basis for applying chemical modifications. These included a modified dimer at position 3 and a monomer at position 7 to represent unmodified and chemically modified siRNA strands. Molecular dynamics simulations and structural modeling were performed to refine these structures through energy minimizations and trajectory calculations, ultimately creating 30,000 unique modeled structures. Features derived from these structures were used to evaluate nine different modeling approaches, detailed in the subsequent sections. Initial training in TensorFlow produced mixed results; however, further optimization using Autogluon,^22^ under several conditions, yielded improved metrics. Notably, the multi-model ensemble approach demonstrated the best performance for datasets without structural modeling in Sets 2, 3, and 4 (Figure 1).

**Figure 1.**
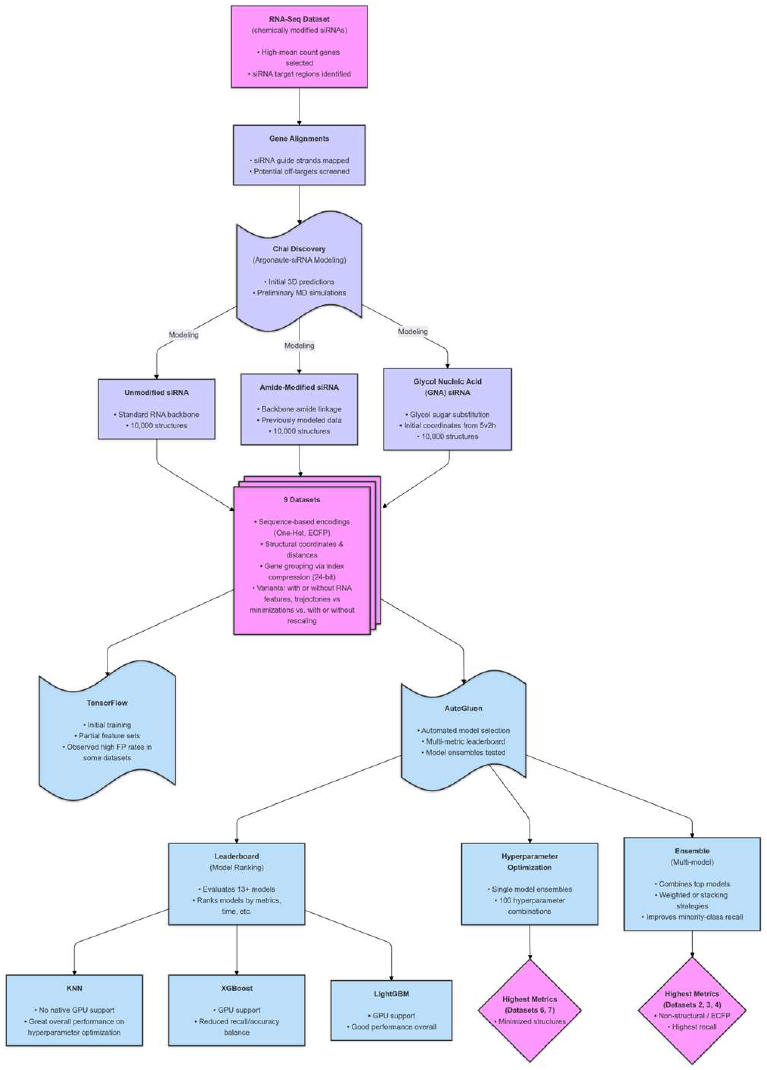
Workflow of the experimental setup. Workflow consisted of 3 phases: experimental, bioinformatics and machine learning. The experimental phase consisted of the production of chemical modified siRNAs and RNA-Seq data. The bioinformatics phase involved aligning the results, structural predictions, molecular modeling and dynamics, to then represent the data in 9 different ways. The machine learning involved initial testing using TensorFlow, and several optimization rounds with Autogluon.

**Figure 2.**
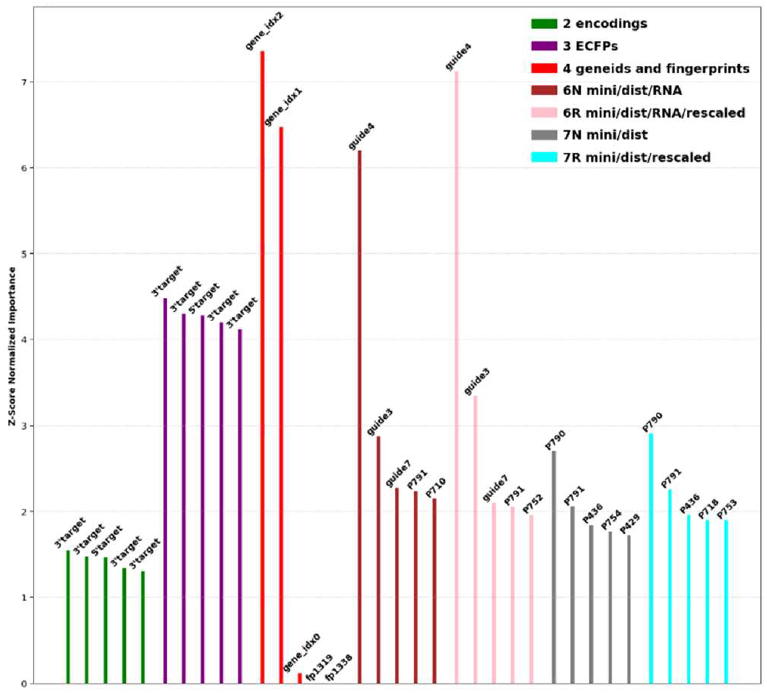
Top 5 features per dataset. Feature importance was assessed using RandomForestClassifier (n_estimators=75). Feature importance was ranked using mean decrease in impurity (Z-Score Normalized).

Hybrid datasets combining diverse feature types, such as sequence-derived and contextual features, have demonstrated improved predictive capacity in biological modeling, underscoring the importance of integrating complementary data sources into feature extraction workflows. Building on this concept, we designed nine unique datasets, each representing chemically modified siRNAs, changes in the Argonaute protein during RISC activation, and the targeted genes (Table 1). Eight of the nine datasets incorporate novel approaches, with Dataset 2 serving as a control. This strategy combines the strengths of molecular dynamics and structural refinement with the interpretability of conventional encodings. By adding detailed spatial and chemical insights to the feature space, this method marks a significant advancement in feature engineering for siRNA and provides a robust framework for off-target effect prediction.

**Table 1.** Summary of the 9 datasets used in this study. Datasets 0 and 1 used data from molecular dynamic trajectories as features, whereas 6 and 7 used minimizations. Dataset 2 used conventional encodings, 3 used a complete set of ECFP fingerprints to represent both the unmodified and modified siRNA nucleobases, and 4 used only key fingerprints along with gene clustering information.

| Dataset | Description | Key Features | Features | Reference |
| --- | --- | --- | --- | --- |
| 0 | Distances calculated after modeling siRNA structures with chemical modifications. | 2 molecular fingerprints, which contain the dinucleotide at position 3-4 and a monomer representing position 7. Distances after trajectories. | 897 | 4f3t |
| 1 | After modeling, full xyz coordinates of each residue were stored and used as features. | Includes coordinates of all protein and RNA residues. No fingerprints included. Used trajectories. | 2638 | 4f3t |
| 2 | Simple RNA encoding of guide strand and predicted target with numerical modification indicators. | Encodings: A = 1, C = 2, G = 3, U = 4, dimer = 5-6, monomer = 7. | 43 | None |
| 3 | Extended Connectivity Fingerprints (ECFPs) based on guide strand and target sequences to give molecular fingerprints. | Fingerprints calculated instead of numerical encoding (as in Dataset 2). | 1345 | None |
| 4 | Uses gene index data as features, and same fingerprints as 0. | Ensembl IDs type-casted into uint8 to give 3 data points (gene index 0-2), same molecular fingerprints as 0. | 100 | None |
| 6N | Distances from minimized structures, used pre-modeling structure as reference. | Distances calculated from the pre-modeling base structure. RNA distances included. | 879 | Base Structure |
| 6R | Same as 6N but with rescaled data (0-255) using range normalization for each feature. | Rescaled distance data for improved comparability across features. | 842 | Base Structure |
| 7N | Distances from minimized structures, used literature reference. | Distances calculated from literature reference, no RNA distances. | 842 | 4f3t |
| 7R | Same as 7N but with rescaled data. | Rescaled distance data for improved comparability across features. | 837 | 4f3t |

The key features generated in this process included fingerprints for chemical modifications, changes in residue-to-residue distances, residue coordinates, as well as gene indices to help group the data. The total number of features varied broadly, from 43 for a simple encoding to over 2600 when extracting full xyz coordinates for each protein and RNA residue. To maintain consistency and reproducibility, the post-modeling structures were aligned to a reference, which was either the experimental structure, 4f3t,^23^ or the base structure obtained from Chai Discovery. Since 4f3t lacks many RNA residues, distances for siRNAs were not calculated in datasets 0, 7N, and 7R. By using coordinates or the base structure, RNA data was obtained for datasets 1, 6N, and 6R. Recognizing challenges with trajectory reproducibility, we introduced minimized-only datasets (6 and 7) and added rescaling to improve feature consistency.

## Results

### Initial Trials

Initial trials compared the performance of feature sets using a 1DCNN model implemented in TensorFlow.^16^ Among the tested datasets, Dataset 0 achieved the highest average precision score. While all trained datasets exhibited similar performance ranges, Dataset 0 consistently outperformed the others. However, the differences in precision across the datasets were modest, suggesting comparable efficacy of the initial feature sets.

Next, we optimized the criteria and evaluated the true positives, false positives and accuracy for Dataset 0. Unfortunately, higher true positive counts (TP > 500) corresponded to lower accuracy (0.4562–0.5667), whereas low true positives (TP ≤ 10) yielded higher accuracy (0.7106–0.7112). This suggests a model bias toward zero outputs, inflating accuracy when true positives are underpredicted. Refinements were needed to balance accuracy and true positive counts. Using Dataset 4, precision improved up to 0.50, and accuracy reached 73%, showing better performance in traditional metrics.

### Autogluon

To further enhance model performance and optimize feature utilization, we employed AutoGluon, a robust automated machine learning framework.^22^ AutoGluon simplifies the process of identifying optimal models by leveraging its leaderboard function to rank model candidates based on their performance. This enabled efficient selection of the best model for each dataset, followed by customized training. AutoGluon also excels at handling diverse data types, automatically tuning hyperparameters, and prioritizing various metrics for improved interpretability and predictive power. Table 2 summarizes the results of models selected via AutoGluon’s leaderboard function and subsequent training. Across datasets (excluding Datasets 0 and 1), true positives (TP) ranged from 304 to 497, false positives (FP) from 175 to 267, precision from 0.61 to 0.67, accuracy from 74% to 78%, and TP/FP ratios from 1.59 to 2.01. Dataset 6R with LightGBM achieving the best overall TP/FP ratio of ~2.

**Table 2.** Initial metric scores using Autogluon.

| Dataset | Model | TP | FP | Precision | Accuracy | TP/FP Ratio |
| --- | --- | --- | --- | --- | --- | --- |
| 0 | LightGBM | 0 | 0 | 0 | 71 | N/A |
| 1 | LightGBM | 72 | 46 | 0.61 | 72 | 1.57 |
| 2 | LightGBM | 497 | 267 | 0.65 | 78 | 1.86 |
| 4 | CatBoost | 471 | 241 | 0.66 | 78 | 1.95 |
| 4 | KNeighbors | 430 | 229 | 0.65 | 77 | 1.88 |
| 4 | XGBoost | 402 | 204 | 0.66 | 77 | 1.97 |
| 6N | LightGBM | 304 | 191 | 0.61 | 74 | 1.59 |
| 6R | LightGBM | 351 | 175 | 0.67 | 76 | 2.01 |
| 7N | LightGBM | 361 | 193 | 0.65 | 76 | 1.86 |
| 7R | LightGBM | 355 | 196 | 0.64 | 76 | 1.81 |

### Feature Importance Analysis

To evaluate the critical features driving siRNA efficacy and off-target prediction, feature importance scores were calculated across all datasets using machine learning models. These scores highlight the most influential properties, including chemical fingerprints, residue-to-residue distances, and protein regions critical for siRNA– Argonaute interactions. Figure 1 summarizes the results, showcasing the relative importance of features across nine datasets. The analysis underscores key trends in how structural and contextual features impact model predictions and provides valuable insights for refining feature generation.

Dataset 2 provides a numeric representation of the siRNA/target duplex, while dataset 3 encoded chemical fingerprints of the same information. In both datasets, the feature importance algorithm identified the ends of the target sequence as the most influential features. These ends tend to be overhanging, exhibiting greater structural flexibility compared to the central regions which will be stabilized by several interactions. Duplexes with higher instability (from mismatches) are likely to experience even greater fluctuations at the terminal positions, and the model was likely able to correlate these regions to the experimental activity levels.

Dataset 4 revealed that the last two gene index values were highly significant due to the sequential assignment of Ensembl identifiers, which inherently grouped related genes. Encoding these identifiers as three uint8 values allowed the higher-order bytes to capture numerical similarities, effectively clustering genes with similar sequences. This explains their strong influence during machine learning training. This finding demonstrates how a simple encoding strategy can inherently group related genes (Figure 3A).

**Figure 3.**
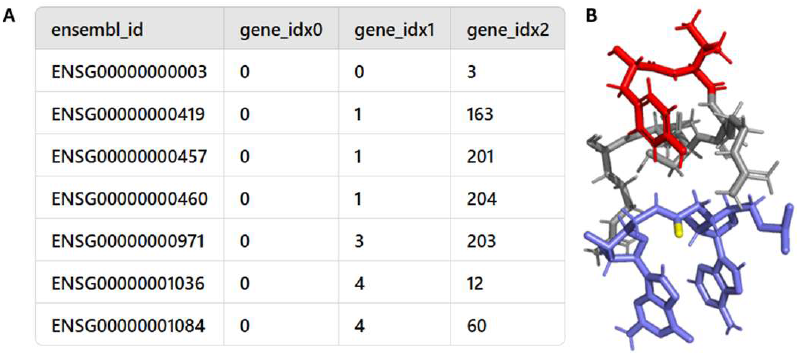
Gene Clustering via 24-bit Identifier Encoding.

Datasets 6 and 7 contained structural features derived from modeling and minimizations. The results highlighted protein residues 790 and 791, which are marked in red in Figure 3B. Feature importance analysis identified these residues as key determinants, as they interact directly with the chemically modified backbone between positions 3 and 4, shown in light blue with the backbone carbonyl highlighted in yellow. Notably, the RNA residues were also the most highly ranked features in datasets 6N and 6R, standing out above all others in the feature importance analysis, reinforcing their central role in RNA interaction and structural stability.

### Hyperparameter Optimization

To further enhance model performance, we utilized AutoGluon’s Hyperparameter Optimization (HPO) framework, which efficiently automates the process of tuning hyperparameters to achieve optimal configurations for machine learning models. By leveraging AutoGluon’s capabilities, the approach ensures systematic exploration of the parameter space, streamlining model development and maximizing predictive performance across multiple metrics, including accuracy, precision, recall, specificity, and F1-score (Figure 4).

**Figure 4.**
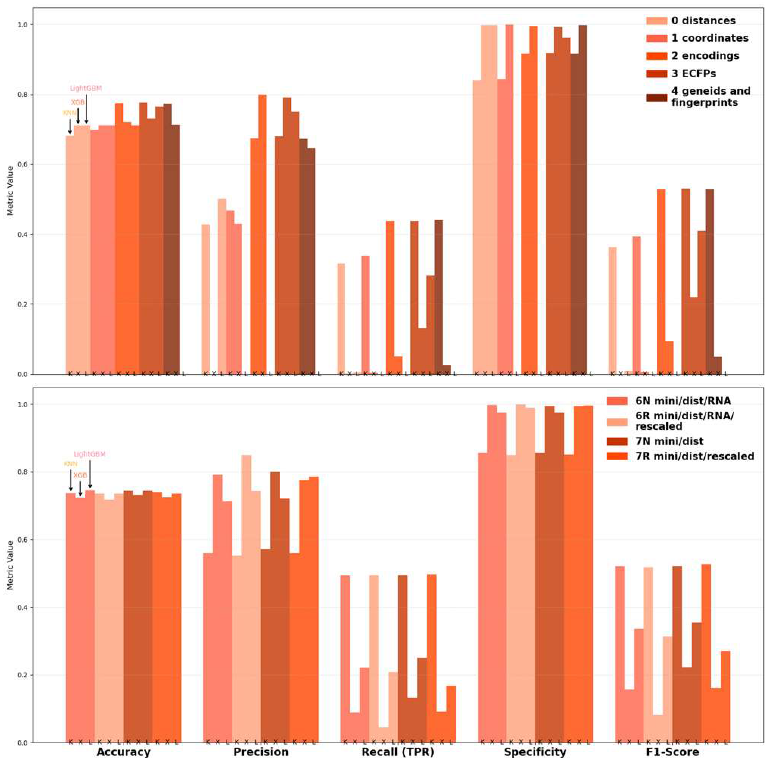
Best Scoring Metric by Model and Dataset. Model order KNN, XGB, LightGBM. Hyperparameter optimization using AutoGluon, showing best score from 10 stratified k-fold cross-validation folds. Performance comparison across five metrics (Accuracy, Precision, Recall, Specificity, F1-Score) for KNN (K), XGBoost (X), and LightGBM (L). Bars are grouped by metric, with models labeled below each group in order. Missing bars indicate that a model received a score of 0, primarily when true positives or true negatives were 0.

**Figure 5.**
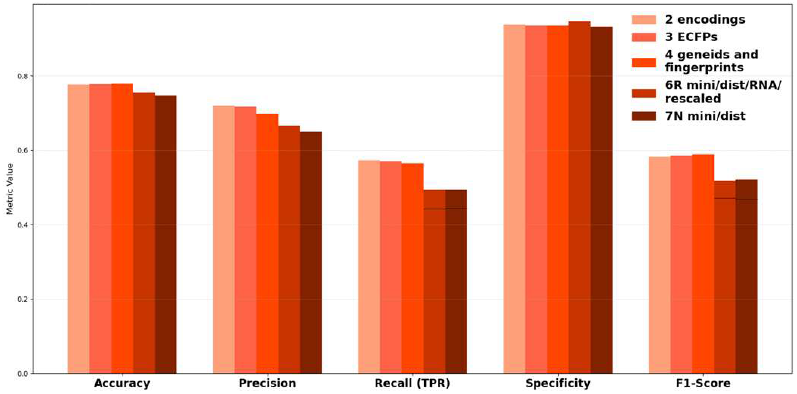
Metric optimization with reduced parameters of the top 5 selected datasets. Multi-model ensembles were used, and 25 different metrics were tested to optimize prediction of the minority class. The top 5 datasets were selected including the simple encodings, fingerprints describing chemical modifications, a hybrid approach using fingerprints and gene grouping, as well as the top 2 datasets produced from modeling.

For datasets 0-4, accuracy scores consistently exceeded across all models, with precision scores ranging from 0.65 to 0.8 for Datasets 2, 3, and 4. Specificity scores were notably high across the board, surpassing 0.85, indicating strong model performance in identifying true negatives. However, recall and F1-scores showed significant variability, with F1-scores only exceeding 0.5 when using KNN. Inflated specificity scores were observed in cases where true negatives (TNs) were disproportionately low, further complicating the interpretation of performance metrics.

With minimized data, accuracy and specificity remained high across all models, with precision reaching close to 0.9 for XGBoost (XGB) on Dataset 6R. However, this performance was misleading as XGB consistently failed to correctly classify the minority class, as reflected by low recall and F1-scores. In contrast, KNN demonstrated the most balanced performance, achieving recall and F1-scores of approximately 0.5 while maintaining high accuracy and specificity (above 0.85). Precision scores for KNN were also satisfactory, exceeding 0.55, making it the most reliable model for minimized datasets.

These findings underscore the importance of balancing metrics during model evaluation. While certain models like XGBoost excelled in precision and specificity, their inability to address class imbalance limited their practical utility. Conversely, KNN provided more consistent performance across metrics, offering a viable solution for off-target prediction in siRNA studies.

### Metric Optimization

Metric optimization was performed to address the mixed results observed during hyperparameter optimization (HPO). By employing multi-model ensembles and testing 25 different metrics, the focus was placed on improving predictions for the minority class. This approach led to accuracy scores consistently approaching 0.8, though precision scores slightly declined, stabilizing above 0.7. Notably, recall and F1-scores improved significantly for Datasets 2, 3, and 4, which were based on simple encodings, chemical fingerprints, and hybrid feature strategies without structural modeling, reaching nearly 0.6.

For minimized datasets (6R and 7N), however, recall and F1-scores were lower compared to non-minimized datasets, indicating a trade-off between model complexity and generalization. To illustrate this discrepancy, a black line was added to the figure, representing the HPO-optimized results for these datasets as a reference point. This comparison underscores the challenges of balancing performance across diverse datasets and optimization methods. Ultimately, metric optimization demonstrated its value in improving minority class predictions, particularly for non-modeled datasets, while emphasizing the need for dataset-specific approaches to enhance overall siRNA off-target prediction.

### Stack Level and Time Optimization

Following metric optimization, stack-level and time optimization was explored using AutoGluon’s ensemble framework, which integrates base models into meta-models at successive stack levels. Notably, despite representing the same underlying data (Figure 6), Dataset 3—employing molecular fingerprints to convert a 21-base siRNA into 672 features—outperformed Dataset 2, which uses a simpler numeric encoding for RNA bases and chemical modifications. Although both datasets were expected to yield comparable training results, the increased numerical richness of Dataset 3 enabled a more precise characterization of siRNA properties, achieving AUPRC scores nearing 0.8.

**Figure 6.**
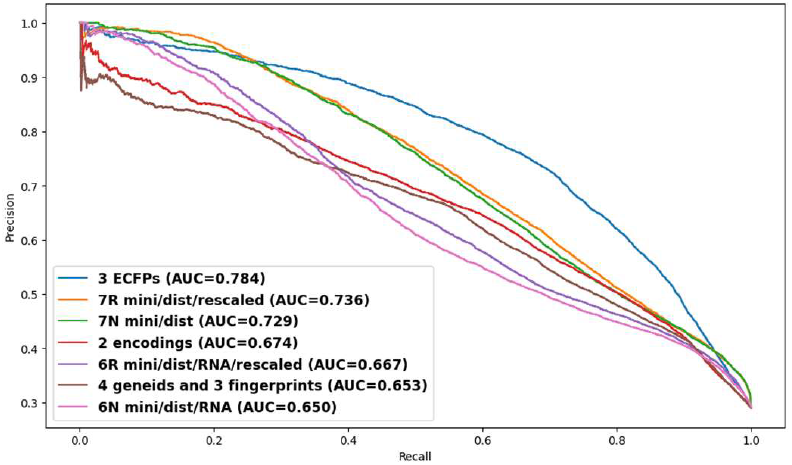
Stack level and time optimization. The 3 ECFPs achieved the highest AUPRC (0.784), while both 7R and 7N performed similarly (~0.73). This highlights the importance of expanding molecular data into full functional groups and leveraging an experimental structure as a reference for improved siRNA off-target predictions.

Given that siRNA off-target prediction remains an undeveloped area in machine learning with limited direct comparisons, our model’s AUPRC scores of 0.784 and 0.736 stand against 0.714 reported for BERT-siRNA24^24^ suggesting differences in how feature encodings and optimization strategies influence precision-recall tradeoffs. While AUPRC is the optimal metric for siRNA off-target prediction, Spearman correlation is often preferred in other analysis, with values of 0.639,^24^ 0.5,^25^ 0.84,^26^ reported across studies, reflecting differences in ranking consistency.

### ECFPs

Figure 7 illustrates the changes in extended connectivity fingerprints (ECFPs) resulting from a progressive replacement of single atoms in a guanosine monomer structure. Starting from an all-carbon base structure, each atom was sequentially replaced with the correct atom, beginning with 1-P, the central phosphorus in the backbone. For example, line 0 represents the fingerprints of the initial all-carbon structure, line 1 shows the results after replacing one carbon with phosphorus (1-P), and line 23 corresponds to the final structure with all substitutions completed.

**Figure 7.**
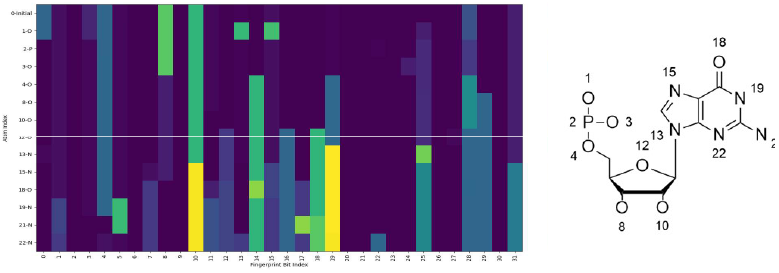
Theoretical demonstration illustrating the features generated by RDKit. Heatmap of ECFP changes in a guanosine monomer during sequential atom substitutions (line 0: all-carbon; line 23: fully substituted). The y-axis shows atomic indices (with atom types) and the x-axis represents fingerprint bit positions, with color intensity reflecting substructure presence. To the right the corresponding atom indices are listed for reference.

The fingerprints were generated using the RDKit library, where each bit in the ECFP encodes local molecular substructures. The y-axis represents atomic indices, labeled alongside their corresponding atom types, while the x-axis corresponds to fingerprint bit positions. Carbon atoms were excluded from visualization since substitutions involving carbon do not alter the fingerprints. The color intensity in the heatmap indicates the fingerprint value at each bit index. ECFPs are generated by iteratively hashing atomic environments into unique numerical identifiers. Each atom is assigned an initial feature based on its properties (e.g., atomic number, hybridization, formal charge, and bond order), with neighboring atoms progressively incorporated up to a defined radius (radius = 1 in this study). These hashed substructures are folded into a fixed-length binary vector (256 bits), encoding the presence or absence of specific patterns. To enhance storage efficiency and facilitate visualization, the binary fingerprints were compressed into unsigned 8-bit integers using numpy’s np.packbits function, preserving the substructural information in a compact form. Significant changes in the fingerprints were observed upon insertion of functional groups. For example, 12-O completes the sugar ring moiety, and the addition of the final nitrogen 22-N completes the nitrogenous base, so these show many signature changes.

This fingerprinting method provides a powerful way to describe chemical modifications by capturing molecular changes across the structure. However, it comes with limitations, such as increased memory usage due to the high dimensionality of the features. Despite its complexity, the method offers valuable insights into the impact of atomic substitutions on molecular properties.

In addition to the scores with dataset 3, notable AUPRC scores were also obtained for both versions of dataset 7. This data was obtained via a process that involved structural prediction and molecular modeling. The unmodified guide-target siRNA/argonaute structure was predicted using Chai Discovery, and then the modification was inserted into the predicted structure using the experimental literature coordinates. Gromacs was then used to setup a water box with ions, and then the complex was minimized up to 3 times to ensure reproducible results. After minimization, distances were calculated as follows: for each minimized structure the solvent and ions were removed, and the siRNA/AGO complex was then aligned to the experimental 4f3t structure. Then for each protein residue, the change of distance was calculated and saved as a feature. 7R provides a couple additional data refinement steps compared to 7N: for each feature, the data was rescaled in the range of 0 to 255 based on the lowest to highest value of the column. With AUPRC scores reaching nearly 0.75, this demonstrates the utility of this method.

### Conclusion

In this work, we explored nine distinct approaches for featurizing chemically modified siRNAs and their off-target interactions, using an RNA-Seq dataset that included 150,000 modified siRNA–gene pairs. Our results demonstrate that computational modeling—particularly structure minimization—provides a powerful framework for generating meaningful 3D-aware features. Although the minimized-structure datasets (notably 7N/7R) performed very well, the top performer was dataset 3, which leveraged extended connectivity fingerprints (ECFPs) derived from the siRNA and intended mRNA target. These findings highlight the importance of capturing both chemical modification details and spatial or energetic properties for robust off-target predictions. Moving forward, we envision integrating the strongest components from multiple feature sets, combining minimized structural data with ECFP-based encodings, for improved predictive performance. Overall, our study underscores the feasibility and promise of using systematic, reproducible structural prediction and modeling approaches—particularly minimization—to guide machine learning models aimed at understanding and mitigating off-target effects in siRNA therapeutics.

### Future Direction

Chai Discovery can incorporate much larger mRNA target strands into the structures, and integrating this capability into the next version could offer deeper insights into siRNA– target interactions, particularly regarding loading or positional effects on mRNA. New features could be selected from highlighted regions, such as those in Figure 3B, placing emphasis on high-impact Argonaute residues (e.g., near positions 790–791). In addition, using a larger dataset will improve predictive performance and could be expanded to include a broader range of chemical modifications, enhancing the model’s ability to detect diverse off-target patterns. Reproducibility in modeling and simulations is essential, and we intend to leverage software that facilitates reliable and repeatable results. Ultimately, the combination of richer datasets, more targeted structural modeling, and reproducible computational workflows will lay the groundwork for a next-generation siRNA design platform that more accurately predicts and mitigates off-target effects.

## Materials and Methods

### Reproducibility

Reproducibility is the most important aspect in experimental data and can often be overlooked when making structural predictions or conducting modeling. To ensure that all data gathered in this study was reproducible, we developed a criteria to compare two or more structures generated from the same starting point. Duplicate structures were aligned to reference 4f3t and distances were found for each protein residue. The relative standard deviations were then calculated for residue within each structure and were then summed. Duplicate structures were then meticulously generated for Chai Discovery, post minimization, and post trajectory, and the results were 3.39, 0.03, and 6.27 respectively. Because of the increased value for trajectories, the time step was reduced to the final settings as listed below to give a ∑RSD of 3.01.

### Minimizations and trajectories

For the amide modified backbone, the partial charges and initial structure were generated as previously described.^27^ For the glycol nucleic acid, the initial coordinates were taken from the published structure 5v2h^19^ and the partial charges were derived using the restrained electrostatic potential approach based on an electrostatic potential grid calculated on the GNAU monomer at HF/6-31G* using NWChem. To incorporate the glycol nucleic acid (GNA), which lacks a conventional sugar moiety and thus precludes the fitting approach used for amide modifications, an alternative strategy was employed. The GNA base, along with its two adjacent nucleotides, was derived from the coordinates of the experimental structure. These adjacent bases served as anchor points for fitting to the corresponding bases in the predicted siRNA–Ago2 complex structure. GROMACS^28^ was used to generate a cubic solvation box ensuring a minimum distance of 1.0 nm between the solute and box edges. The protein/nucleic acid complexes were surrounded with TIP3P water molecules and ions were added to neutralize the system. Molecular dynamics and minimizations were carried out with AMBER 99 (ff99SB).

### Chai Discovery

Predicted structures of siRNA duplexes within the human Argonaute-2 protein were generated using Chai-Lab v0.0.1 with the following parameters: num_trunk_recycles=3, num_diffn_timesteps=200, seed=42, and use_esm_embeddings=True. The highest-scoring prediction was retained.

### Conversion of Ensembl Gene Identifiers to 24-bit Unsigned Integer Representation

Ensembl gene identifiers were compressed into a 24-bit unsigned integer representation to enable efficient storage and processing. Each identifier was parsed to remove the prefix ENSG and any leading zeros, leaving only the numeric portion. The numeric portion was then split into three unsigned 8-bit integers (uint8), corresponding to the most significant byte, middle byte, and least significant byte, using bitwise operations. Specifically, the most significant byte was extracted as (n >> 16) & 0xFF, the middle byte as (n >> 8) & 0xFF, and the least significant byte as n & 0xFF, where n is the numeric portion of the identifier. These three uint8 values were stored as separate columns in the dataset, labeled gene_idx0, gene_idx1, and gene_idx2, to facilitate downstream analyses. This approach ensures the preservation of uniqueness in the compressed format while minimizing storage requirements.

### ECFPs

For datasets 0 and 4, the fingerprints were generated for dimers at position 3 and 4, then for monomers at position 7. The fingerprint of the dimer was made by selecting the unmodified AG or amide modified AG, and creating a fingerprint with nBits=512, radius=2. The fingerprint of the monomer was made by selecting either the unmodified U or modified GNA U at position 7, and creating a fingerprint nBits=256, radius=1. The binary vectors were compressed into unsigned 8-bit integers using numpy’s np.packbits, making 64 and 32 features, respectively.

For dataset 3, a different approach was used. To focus on functional groups, the hydrogens were first removed. Instead of using dimers, every position was treated as a monomer, selecting only the atoms associated with that position, as they were assigned within Amber. And the above settings for monomers were used, producing 32 features per position.

### Autogluon

Hyperparameter optimization was conducted using AutoGluon’s hyperparameter_tune_kwargs function with varied trial numbers. This process targeted key models including LightGBM, XGBoost, CatBoost, and KNN. Hyperparameter ranges were defined according to each model’s key parameters and tested across multiple predefined configurations. To assess the value of additional trials, the number of optimization iterations was systematically varied. Results indicated that increasing the number of trials beyond two did not consistently improve performance, suggesting diminishing returns from extended optimization.

Metric optimization was set up by removing skf and varying the seed. All the available metrics were tested using the default ensembl with all available models. balanced_accuracy, roc_auc_ovo_macro, recall, and f1_weighted were selected and used to train all the datasets with 10 different seed splits.

Stack level optimization was performed with metric roc_auc_ovo_macro. The num_stack_levels and time_limit were varied from 4 to 10 and from 130 to 900 s respectively. The performance of various feature sets was evaluated using precision-recall curves.

## Dataset Availability

To ensure transparency, the code and the in-house RNA-Seq dataset used in this study are available at https://github.com/mrichter0/siRNA-Features

## Notes

https://github.com/mrichter0/siRNAFeatures

### Competing Interest Statement

The authors have declared no competing interest.

